# scMOBA: A conversational single-cell Multi-Omics Brain Agent across species

**DOI:** 10.64898/2025.12.01.691565

**Authors:** Ran Wei, Ziyao Zhang, Jianle Sun, Yongkang Sun, Juan Meng, Peng Zheng, Chaoqi Liang, Fanyi Meng, Wanli Ouyang, Lei Bai, Peng Ye, Yidi Sun

**Affiliations:** Shanghai Artificial Intelligence Laboratory, Shanghai, China; Institute of Neuroscience, CAS Center for Excellence in Brain Science and Intelligence Technology, Chinese Academy of Sciences, Shanghai 200031, China; University of the Chinese Academy of Sciences, Beijing 100049, China; Carnegie Mellon University, Pittsburgh 15213, PA, US; University of Trento, Trento 38122, TN, Italy; Harbin Institute of Technology, Harbin, China; The Chinese University of Hong Kong, Hong Kong, China; State Key Laboratory of Genetic Evolution & Animal Models, Shanghai, 200031, China; Shanghai Key Laboratory of Precision Gene Editing and Clinical Translation, Shanghai, 200031, China

## Abstract

Single-cell and spatial multi-omics are revolutionizing our understanding of the complexity in the developmental, aging, and diseased brain, but integrating this knowledge across modalities and species remains challenging. To bridge this gap, we propose scMOBA, a conversational single-cell Multi-Omics Brain Agent established by a large language model, a gene encoder, and a cross-attention projector. The scMOBA was pre-trained on 130 million single-cell and spatial multi-omics data spanning the entire brain across diverse species, development, aging and diseases. The pre-training utilized a novel multi-omics Feature-Question-Answer (FQA) paradigm, enabling the model to generate biological answers from feature inputs and textual queries. This unique scheme facilitates superior zero-shot inference capabilities without requiring additional fine-tuning. We demonstrate that scMOBA achieves state-of-the-art performance in fine-grained cell type classification across different species and modalities, as well as in batch correction and multi-omics data integration. Furthermore, scMOBA significantly boosts the accuracy of kinds of critical downstream tasks, including cell-type specific aging clock construction and disease status prediction. Overall, scMOBA serves as a powerful scientific discovery engine for multi-omics brain research, advancing the precision prediction and early intervention of neurological aging and diseases.

## Introduction

A comprehensive understanding of the cellular and molecular architecture of the brain forms the foundation for deciphering the neural mechanisms underlying cognition, behavior, and disease. Recent advances in single-cell and spatial multi-omics have accelerated the understanding of cellular composition and their organization principles in the brain of Drosophila, zebrafish, mouse and primate brains^1–10^. Datasets encompassing different developmental stages and brain disorders have begun to reveal certain cell types and molecular features associated with development, aging and disease progression^11–17^. However, most existing studies were restricted to single modality, limited regions, specific species or a certain trait, resulting in significant gaps in our understanding of cell type consensus and the complex interplay across molecular signatures from genomics, epigenomics, transcriptomics and proteomics data. The integration of large-scale published datasets may provide a more complete landscape of the brain cells, enabling cross-regional, cross-modal, and cross-species comparisons to pinpoint key cell populations in neuronal processes and neurodegenerative diseases.

Despite the challenge posed in the diversity of omics methodologies and the inherent heterogeneity of datasets, the success of large-scale, self-supervised foundation models presents a significant opportunity for the integration. Pre-trained models such as scGPT^18^, Geneformer^19^, scFoundation^20^ and GeneCompass^21^ utilizing single-cell transcriptome data have demonstrated capability across a broad range of downstream tasks including cell clustering, cell type annotation and gene perturbation simulation. Likewise, general expression transformer (GET)^22^ has been introduced to uncover transcriptional regulation based on chromatin accessibility data. Nicheformer^23^, CellPLM^24^ and BrainBacon^25^ integrating spatial data enable the prediction of the spatial context of dissociated cells. Moreover, recent advance like SCARF^26^ has begun to expand beyond the single-cell transcriptome foundation model to single cell ATAC-seq and RNA-seq foundation model. Nephrobase Cell+^27^, a multi-modal single-cell foundation model has been developed for decoding the kidney. Nevertheless, a foundation model for the integrative analysis of multi-omics data spanning diverse brain regions, developmental stages, aging, and disease conditions across multiple species is still lacking.

In addition, a critical gap exists between powerful foundation models and the scientists who could benefit from them, as the current foundation models typically require fine-tuning to each specific task to achieve accurate representations and predictions. However, most researchers in the field of neuroscience do not possess the coding skills required for such implementations, limiting their application to computational experts in specific fields. This issue is exacerbated because these foundation models are not user-friendly or conversational. To address this gap, we propose a natural language interface capable of executing complex tasks from straightforward English commands. The proven success of intuitive, instruction-following large language models (LLMs) like ChatGPT^28^, LLaMA^29^ and DeepSeek^30^, which are reshaping industries, provides a powerful validation for this approach. We thus envision the natural progression towards the introduction of biological expert systems that offer similar conversational capabilities in the field of single-cell and spatial multi-omics brain data.

Actually, recent studies have attempted to use LLMs to generate gene embeddings from textual descriptions of individual genes for scRNA-seq analysis^31^. However, this framework converts scRNA-seq expression profiles into a textual format (selecting the top-K highly expressed genes and arranging their gene names), the inherent hallucination problem of LLMs raises concerns about the reliability and reproducibility of biologically meaningful representations and predictions. Moreover, such a lossy text-based representation is fundamentally unable to capture the dynamic and context-dependent roles of genetic features and cells across different biological states. Inspired by visual–language models (VLMs) that jointly process images and text^32–34^, we thus proposed to develop a multimodal framework capable of integrating multi-omics data with natural language. By treating feature abundance profiles from multi-omics data as analogous to visual data and representing biological states and insights in natural language, the model can take both feature abundance and text describing biological information as inputs, enabling multimodal understanding. We hypothesize that this multimodal framework can jointly model data from diverse omics modalities and multiple species, enabling their alignment within a unified text-informed representation space.

Thus, in this study, we proposed a conversational <u>s</u>ingle-<u>c</u>ell <u>M</u>ulti-<u>O</u>mics <u>B</u>rain <u>A</u>gent across species named scMOBA, which allows an integrated comprehension of cross-species and multi-omics single-cell data in the brain. The scMOBA architecture consists of three key components: a gene encoder, a cross-attention projector and a large language model. Leveraging natural language processing, scMOBA was pre-trained on a brain corpus of 130 million single-cell and spatial multi-omics profiles, covering whole brain across various species and biological conditions. By adopting a multi-omics feature–question–answer (FQA) paradigm, the model learned to generate biological responses directly from feature abundance inputs and textual queries. Since this training scheme aligns with real downstream applications, scMOBA demonstrates zero-shot inference capabilities without the need for additional fine-tuning, including accurate prediction for fine-grained cell type classification in datasets from different species or modality, aging clock construction in various cell types and prediction of species-specific or disease-specific cell types. In addition, scMOBA demonstrated the state-of-art performance in batch correction, multi-omics and cross-species data integration. Overall, scMOBA represented a conversational multi-omics brain agent for empowering major brain atlas initiatives by providing a powerful scientific discovery engine for multi-omics research and advancing the precision prediction and early intervention of aging and diseases in the brain.

## Results

### The architecture of scMOBA and FQA pretraining

To develop an intelligent agent for cross-species multi-omics data integration and mapping in the brain, we first constructed a large-scale comprehensive brain corpus comprising 130 million single-cell multi-omics data spanning evolutionary distance, molecular levels, various neuroanatomical regions, developmental stages and pathological brains for pre-training (Fig. 1a and Supplementary Table 1). Specifically, we included snRNA-seq, snATAC-seq and spatial transcriptome data from each of the four mammalian species (human, macaque, marmoset and mouse), covering over 124 cortical and subcortical brain regions, the entire lifespan (embryonic, infancy, adolescent, adult and aged), and various brain diseases (e.g., psychiatric, cognitive impairment, and neurodegeneration) (Extended Data Fig. 1a, b). To ensure the consistency of metadata from the large-scale and diverse high-dimensional datasets, we applied an LLM-assisted manual curation procedure to align the cell type taxonomy, brain region origins, developmental stages and disease status from different datasets (Extended Data Fig. 1c and Supplementary Table 2). Based on the curated metadata, we designed a number of 225 million feature-question–answering pairs covering major cell properties of our brain corpus for subsequently training multi-modal large language models (Fig. 1b).

**Fig. 1.**
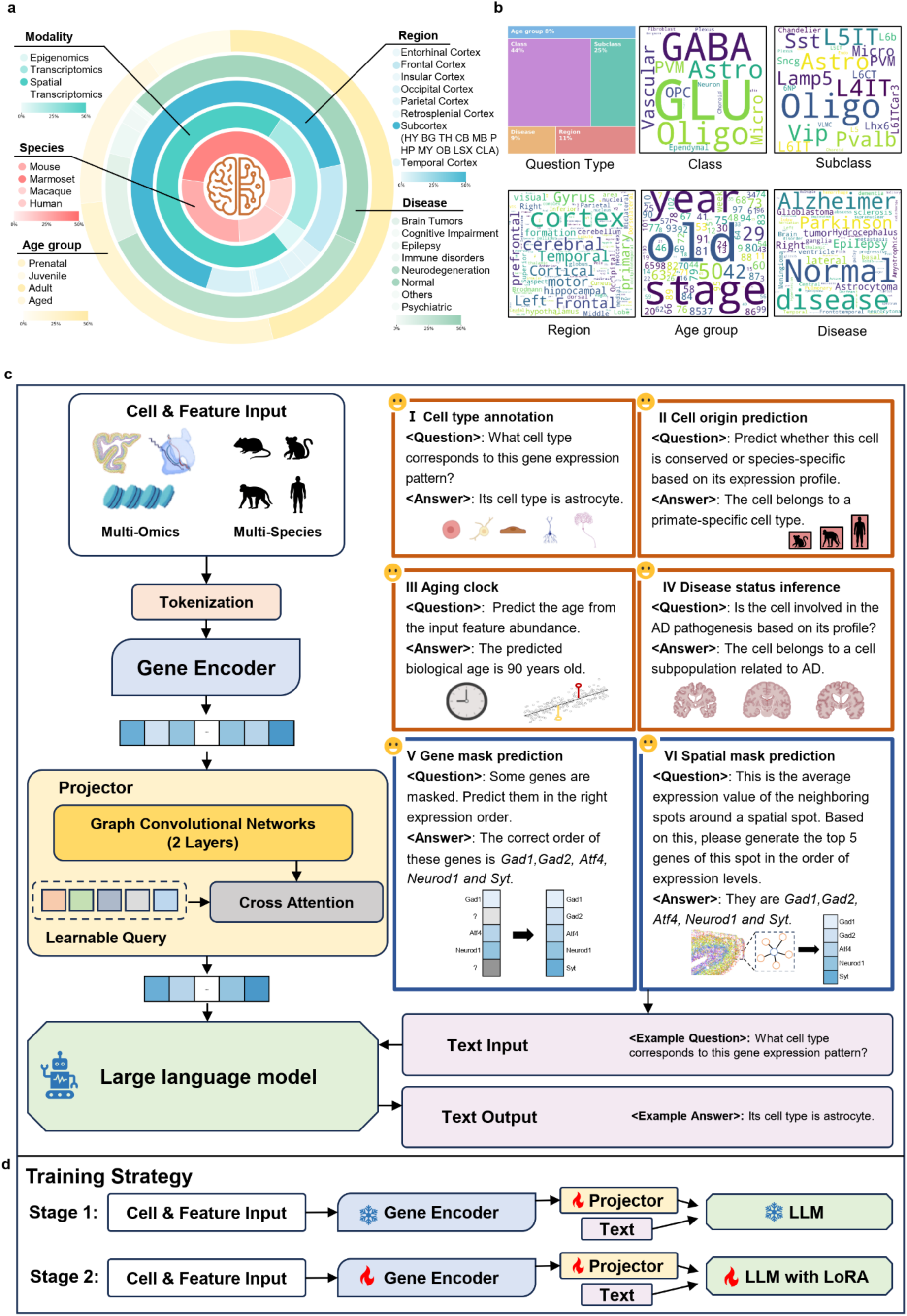
Overview of scMOBA. **a**, Summary of the pre-training datasets, comprising approximately 130 million single-cell multi-omics data from various brain regions across four species and diverse developmental stages and disease status. **b**, Word clouds summarizing the distribution of question types, cell classes, cell subclasses, brain regions, ages and disease types of the text used for constructing the question-answer pairs. **c**, Schematic overview of scMOBA architecture. **Left**, The architecture consists of three major components: a gene encoder, which converts gene expression or activity matrix into gene and cell embeddings; a projector, which model gene–gene interactions based on the encoded gene representations and extract the most relevant features for downstream textual reasoning to the language model; and a large language model, which receives the concatenated gene-derived features and text-formatted queries to generate natural language outputs. Right, Six kinds of question-answer pair types constructed in the pre-training. **d**, Training strategy of scMOBA. Parameters of both the gene encoder and the language model are frozen in the first stage, while all components are active for training in the second stage.

The core design principle of scMOBA lied in bridging the semantic gap between the language of multi-omics data and the language of humans. To interpret complex multi-omics brain datasets, scMOBA applied a feature–question–answering framework that integrates a gene encoder, a projector, and a LLM (Fig. 1c and Extended Data Fig. 1d). The architecture began by processing tokenized cell and feature inputs (e.g., gene expression or snATAC-seq-derived gene activity score) through a dedicated gene encoder. The resulting embeddings were then passed through a projector that extracts key genetic features and maps them into the language space of the LLM. The LLM received and concatenated both the feature abundance representations and the corresponding textual questions as input, and generates biological answers in natural language. The scMOBA processed each omics data separately, and employed a unified feature–question–answer paradigm to handle a wide spectrum of tasks (from cell type annotation to disease status prediction), where each supervised task was formulated as a question with a natural language answer. The architecture of scMOBA offered flexible downstream tasks suitable for facile cell type annotation, cell origin prediction, aging clock construction and disease status inference. In addition, the unsupervised strategy of randomly masking tokens and reconstructing the missing signals to capture context-specific patterns was also incorporated into this unified paradigm (Fig. 1c). With the feature–question–answer design, it enabled the integration of supervised and unsupervised signals into a unified training framework, while allowing data from different modalities and species to be represented under the same formulation.

In this framework, scMOBA was trained to generate accurate textual answers given feature abundance data and natural-language questions. Since both the pre-training and downstream tasks share the same feature–question–answer formulation, their optimization objectives are fully aligned. This design ensured that scMOBA learns a unified objective across pre-training and downstream tasks, distinguishing from existing models that require task-specific architectures and loss functions. To achieve robust cross-modal alignment, scMOBA employed a two-stage pre-training scheme that progressively integrates feature abundance representations with the LLM’s semantic space while retaining the pretrained strengths of each modality. In the first stage, only the projector was trained with the other two modules frozen to achieve cross-modal alignment. In the second stage, all parameters were jointly optimized, and the LLM is fine-tuned using Low-Rank Adaptation (LoRA) for efficient multimodal learning (Fig. 1d). This hierarchical training enabled scMOBA to bridge the semantic gap between the language of genetic features and the language of humans. As a result, the model can reason across diverse multi-omics and cross-species datasets for both predictive and generative analyses.

### Fine-grained cell type prediction

To evaluate whether scMOBA captures brain cell identities, we conducted extensive zero-shot cell type annotation task for comprehensive evaluations across multiple independent datasets from disparate omics layers and species that were unseen in the pre-training stage (Fig. 2a). First, we asked scMOBA to predict cell types in a snRNA-seq dataset from the motor cortex of human^35^. Notably, scMOBA achieved over 99% accuracy in predicting the cell classes (Fig. 2b). The cell classes predicted by the scMOBA embeddings shown by UMAP perfectly matched with the original cell classification for glutamatergic neurons (GLU), GABAergic neurons (GABA), astrocytes (Astro), oligodendrocyte (Oligo), oligodendrocyte progenitor cells (OPC), microglia (Micro) and vascular cells using expression profiles of highly variable genes (Extended Data Fig. 2a). For the more refined cell subclass labels, the predicted cell labels by scMOBA showed well coherence with the original labels annotated in the dataset for most of cell subclasses in this independent dataset, with an average accuracy of 93.1% (Fig. 2c, d). A few mix-ups were observed for laminar IT cells and vascular cell types, possibly caused by the high similarity between IT cell types and the limited number of vascular cell types (Fig. 2d). The reliability of cell type annotation was further demonstrated by the canonical marker genes with uniquely top ranks in the corresponding cell types (Extended Data Fig. 2b). For examples, top weighted genes for L2/3 IT were significantly enriched in axonogenesis, neuron projection morphogenesis and regulation synapse organization pathways, and top ranked genes for ASCs included *GFAP*, *SLC1A2* and *S100B* that were involved in regulation of gliogenesis and glial cell differentiation (Extended Data Fig. 2c).

**Fig. 2.**
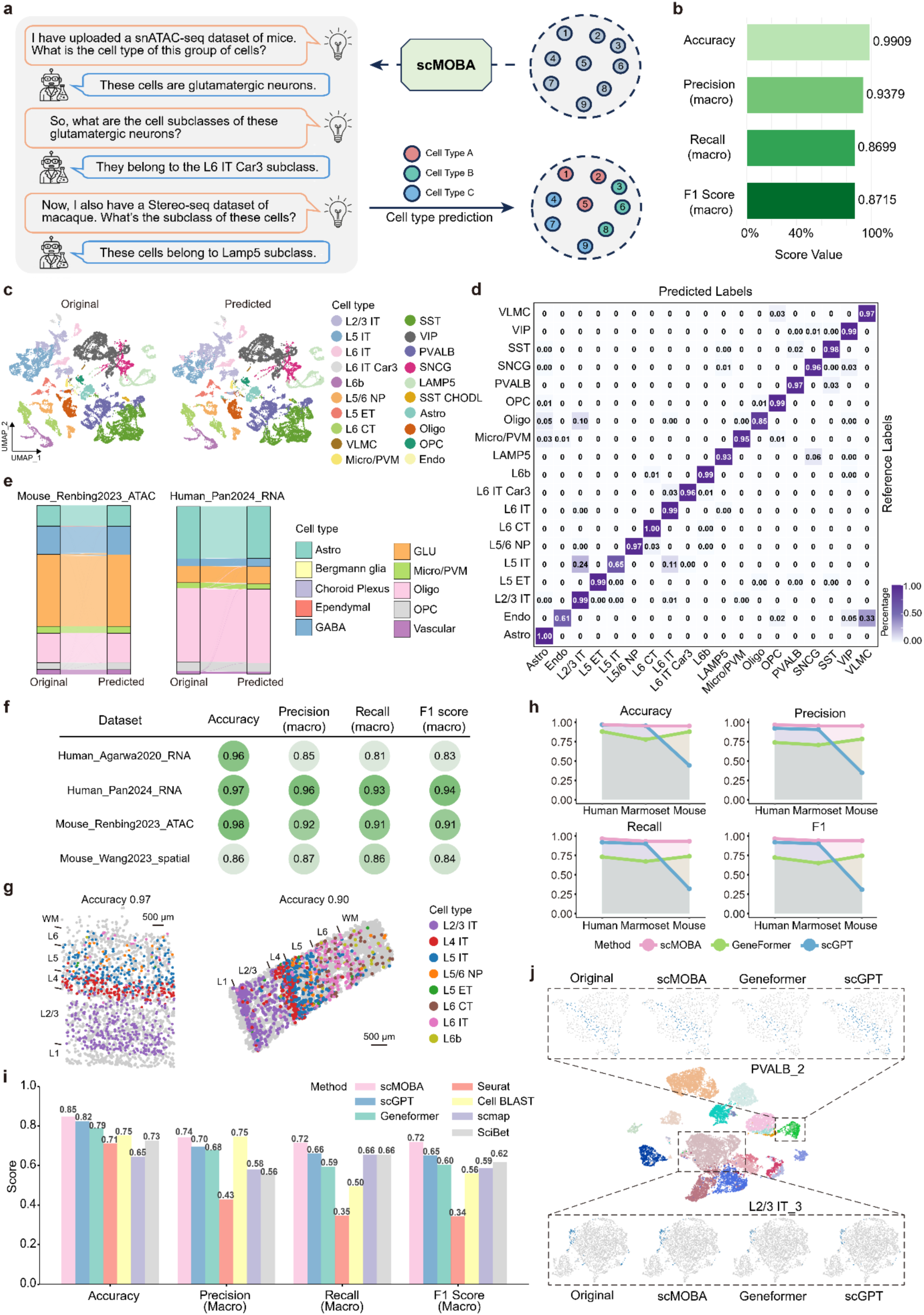
Performance of cell type annotation using scMOBA. **a**, Visualization of QA pairs used for annotation of example cell subclasses. **b**, Evaluation of scMOBA for cell class annotation for an unseen M1 snRNA-seq dataset^35^ at the zero-shot mode. **c**, UMAP plots of gene expression profiles from the M1 snRNA-seq dataset, colored by cell subclasses provided in the original study (left) and those predicted by scMOBA (right). **d**, Confusion matrix heatmap showing the consistency between predicted cell subclasses by scMOBA at the zero-shot mode and originally annotated ones in the M1 dataset. **e**, Sankey plots showing the correspondence of scMOBA-predicted cell types and cell labels provided by the original study in the snATAC-seq^38^ and snRNA-seq^72^ datasets. **f**, Overview of the prediction performance metrics of scMOBA in five independent datasets. **g**, Spatial visualization of predicted cell types by scMOBA in the MERFISH dataset from MTG and STG regions^39^. **h**, Comparison of cell subclass annotation performance of scMOBA, Geneformer and scGPT. Noted that the two other models were not able to predict cell types without fine-tuning, we applied a few-shot strategy using 20% of the dataset as training set and 80% as test set for fair comparison. **i**, Evaluation of the annotation performance for cell clusters using different methods. **j**, UMAP visualizations of example cell clusters predicted by scMOBA, scGPT and Geneformer and cell labels provided by the original study^73^.

We also tested the ability of scMOBA in predicting cell types for query cells from different species or profiled with different omics data types in addition to single-nucleus transcriptome. To achieve this, we retrieved two snRNA-seq datasets from different human brain regions^36,37^, two snRNA-seq datasets from the motor cortex of marmosets and mice^35^, a snATAC-seq dataset from the adult mouse brain^38^, and two spatial transcriptome datasets using MERFISH^39^ and STARmap^40^ techniques. The scMOBA showed consistently high zero-shot prediction accuracy in cell type annotation in the snRNA-seq datasets from humans, marmosets and mice, which were demonstrated by the well consistency in the predicted and the original cell type labels (Fig. 2e, f and Extended Data Fig. 2d). In addition, scMOBA also accurately predicted cell types for the snATAC-seq dataset and the two spatial transcriptome datasets from human and mouse brain (Fig. 2e, f). Specifically, scMOBA predicted cell types showed clear laminar distribution in the spatial transcriptome maps (Fig. 2g). The cross-species and cross-modality abilities of scMOBA for predicting cell labels were further shown by high values in classification metrics of accuracy, precision, recall and macroF1 in the five independent datasets (Fig. 2f, g).

We next benchmarked the ability of scMOBA in cell type annotation with previously published single-cell foundation models, Geneformer^19^ and scGPT^18^. Given that the latter two models require fine-tuning for predicting cell types, we performed a small data training for all the three models in this task. To determine a suitable training–validation split for small-data training, we evaluated the performance of all three models using 10 cutoffs (5% to 50%, in 5% increments) as the training dataset. We found that our model was already able to learn effectively at the 5% split cutoff (Extended Data Fig. 2e). Using this small-data strategy on the snRNA-seq datasets from the primary motor cortex of humans, marmosets, and mice^35^, the results showed that scMOBA outperformed the other models in terms of accuracy, precision, recall, and F1 scores for predicting cell subclasses across all three species (Fig. 2h). Although scGPT showed similar prediction efficacy in the human and marmoset datasets with that of scMOBA, its performance drastically reduced in predicting mouse cell subclasses (Fig. 2h). These results highlight that our model requires only a small number of training samples to achieve reliable cell subclass classification.

Furthermore, we demonstrated the ability of scMOBA for identifying fine-grained cell subtypes, we fine-tuned the model on 80% of the independent dataset and evaluated on the 20% test set for predicting over 100 cell subtypes (see Methods). The fine-tuned model showed well alignment with cell subtypes classified by the original study with an accuracy of 0.85 (Fig. 2i). We also benchmarked the fine-tuned scMOBA with the other two single-cell foundation models and deep learning models developed for cell type classification, and found that scMOBA constantly outperformed the other methods in all classification metrics, including accuracy, precision, recall and F1 (Fig. 2i). The better performance of scMOBA was further visualized by UMAPs showing representative cell subtypes of PVALB neurons and L2/3 IT neurons, where cell annotation predicted by scMOBA showed higher resemblance to the original annotation (Fig. 2j and Extended Data Fig. 2f). These results indicated that scMOBA was particularly well-suited for accurate annotation of fine-grained and even ultra-fine cell subtypes. Together, facilitated by the conversational feature, scMOBA enabled accurate and convenient cell type annotation for brain cells from diverse species and data modalities.

### Multi-modality data integration

Given the remarkable capability of scMOBA in cell type classification across modality data, we next assessed whether its representations could integrate single-cell datasets by separating true biological difference from technical batch effects. To learn unified cell representations, we fine-tuned scMOBA with two different modes: supervised training with cell labels available in the integrated datasets, and semi-supervised training based on pseudolabels from the pretrained model (Fig. 3a). We firstly evaluated the batch removal ability of scMOBA using two human snRNA-seq datasets from different donors^35^. The results showed that scMOBA clearly separated distinct cell types while the same cell types from different batches were integrated together, even in the semi-supervised pseudolabel mode (Fig. 3b). We then conducted a benchmark analysis with two single-cell foundation models - Geneformer and scGPT, and five commonly used multi-modal integration methods - Harmony^41^, Seurat v4 RPCA^42^, Seurat V3 CCA^43^, fastMNN^44^, and scVI^45^. The results indicated that while the other models successfully corrected batch effects across the datasets, their ability to distinguish distinct cell types was comparatively limited next to scMOBA (Fig. 3b and Extended Data Fig. 3a). The superior performance of scMOBA was also demonstrated by the highest batch removal score of 0.99 and biological conservation score 0.91 among all the methods. The total integration score (summation of 60% batch removal score and 40% biological conservation score) of scMOBA achieved 0.94, outperforming the second ranked method by over 20% (Fig. 3c). The robust batch correction ability of scMOBA was further illustrated by the successful integration of different spatial transcriptome sections from the mouse brain^3^ (Extended Data Fig. 3b).

**Fig. 3.**
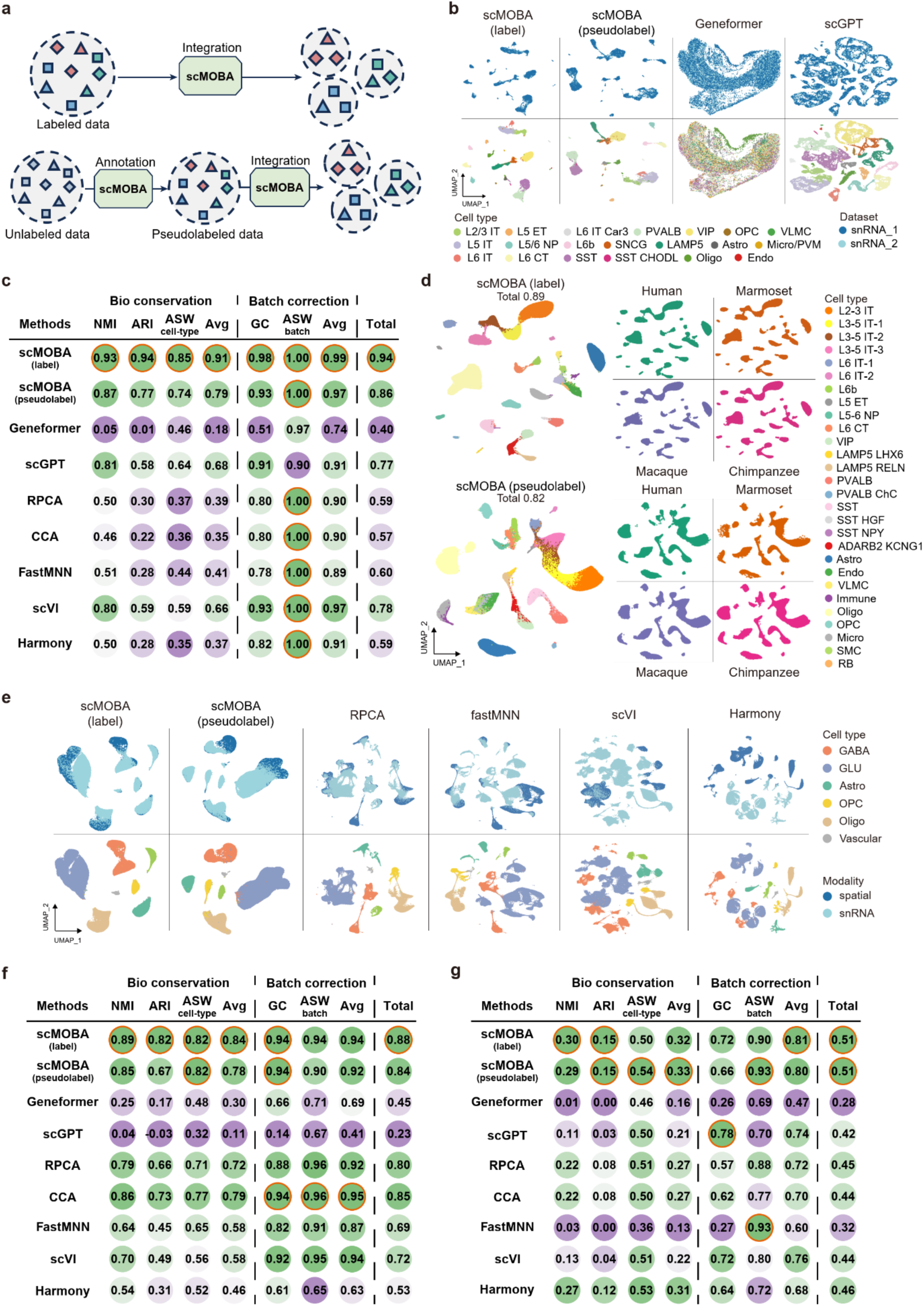
The embeddings of scMOBA result in better integration of multi-omics data from different batches. **a**, Diagram showing the integration of two datasets using the supervised (label) or semi-supervised (pseudo label) fine-tuning strategy of scMOBA. **b**, UMAP plots showing the integration results of two batched snRNA-seq datasets using cell embeddings of two fine-tuning strategies of scMOBA, Geneformer and scGPT. Cells are labeled by their cell types. **c**, Summary of the accuracy results, including metrics of embedding accuracy and cell-alignment accuracy, under different parameter settings of scMOBA. **d**, UMAP visualization of snRNA-seq datasets across four primate species using cell embeddings derived from supervised and semi-supervised fine-tuned scMOBA. The left and right UMAPs were colored by cell types and species, respectively. **e**, UMAP visualization of the tested snRNA-seq and snATAC-seq datasets using cell embeddings derived from fine-tuned scMOBA, Harmony, RPCA, CCA, FastMNN and scVI. **f**, Benchmarking performance of scMOBA in integrating snRNA-seq and snATAC-seq datasets compared with other methods. **g**, Benchmarking performance of scMOBA in integrating snRNA-seq and spatial transcriptome datasets compared with other methods.

Next, we evaluated the ability of scMOBA in integrating cross-species data using snRNA-seq datasets from the primary frontal cortex of human, chimpanzee, macaque and marmoset^8^. Dimensional reduction of scMOBA embeddings showed that the same cell subclasses cluster consistently across the four primate species (Fig. 3d). It was noteworthy that the semi-supervision pseudolabel mode of scMOBA achieved as high as 0.82 in total integration score (Fig. 3d), highlighting the biological relevance of the pretrained embeddings of scMOBA. The superior cross-species integration performance of scMOBA was also supported by the higher biological conservation score and batch correction score over other compared methods (Extended Data Fig. 3c).

In addition, we compared scMOBA with the other seven methods in integrating unpaired snRNA-seq and snATAC-seq data from the primary motor cortex of humans^46,47^. Notably, scMOBA perfectly integrated the two different data modalities while preserving the original cell type information in both supervised and semi-supervised manners, with different cell types separated clearly in the UMAP (Fig. 3e). By contrast, the other two single-cell foundational models could barely integrated snATAC-seq with snRNA-seq modality, probably because of the lack of chromatin accessibility data in the pre-training process (Extended Data Fig. 3d). For other conventional methods, it was noteworthy that Seurat v3 CCA showed much worse biological conservation score (0.79 vs. 0.84), though a slightly higher batch removal capability can be achieved (0.95 versus. 0.94), visible through the better alignment of two data modalities in the UMAPs (Fig. 3e, f).

Finally, we also evaluated the robustness of scMOBA in integrating snRNA-seq and spatial transcriptome data, crucial for assigning cell types from snRNA-seq reference to spatial locations given the limited gene coverage and resolution of spatial transcriptome techniques^48^. We retrieved two independent snRNA-seq and Stereo-seq datasets from the macaque cortex^6^, and fine-tuned the scMOBA for unified cell representations. Similarly, scMOBA achieved superior performance in both batch removal and biological conservation metrics, with at least 10% higher scores than the other benchmarked methods (Fig. 3g). Although cell classes from the two data modalities could be separated after integration for most of these methods, the boundaries of different cell classes were much clearer in the UMAP after scMOBA integration (Extended Data Fig. 3e). Our semi-supervised fine-tuning model presents a distinct advantage for single-cell multi-omics and spatial transcriptomics integration, as it directly leverages the largely unlabeled nature of spatial data to improve cell type identification. Overall, scMOBA demonstrates superior ability in integrating multi-omics data in terms of biological conservation and batch removal metrics.

### Cross-species cell type annotation and comparison

While large-scale cell atlases have broadened our understanding of the brain structure, they are limited by an inability to perform cross-species analysis. Such joint analysis is essential for understanding evolutionary processes, like identifying conserved cell types and the gene programs behind their similarities and differences. Thus, we next assessed the capacity of scMOBA in cell type annotation and comparison across diverse organisms in both zero-shot and fine-tuned manners (Fig. 4a). We started by evaluating the cell type annotation performance of scMOBA in the zero-shot mode using a snRNA-seq dataset collected from the chimpanzee and rhesus macaque dlPFC region^8^. Even though the two species were unseen species in pretraining, scMOBA achieved as high as 0.99 and 1.00 accuracy in predicting the cell identities, respectively (Fig. 4b and Extended Data Fig. 4a). The generalization capacity of scMOBA over the unseen species was further demonstrated by over 99% consistency in the predicted and original cell labels shown by the confusion matrix (Fig. 4c and Extended Data Fig. 4b). In addition, scMOBA distinguished over 20 finer-resolution laminar glutamatergic neurons, CGE and MGE GABAergic neurons, and various glial cell types with an accuracy of 0.87 in chimpanzee and 0.83 in Rhesus macaque without any fine-tuning (Fig. 4d, e and Extended Data Fig. 4c, d), indicating its robustness in cross-species scenarios.

**Fig. 4.**
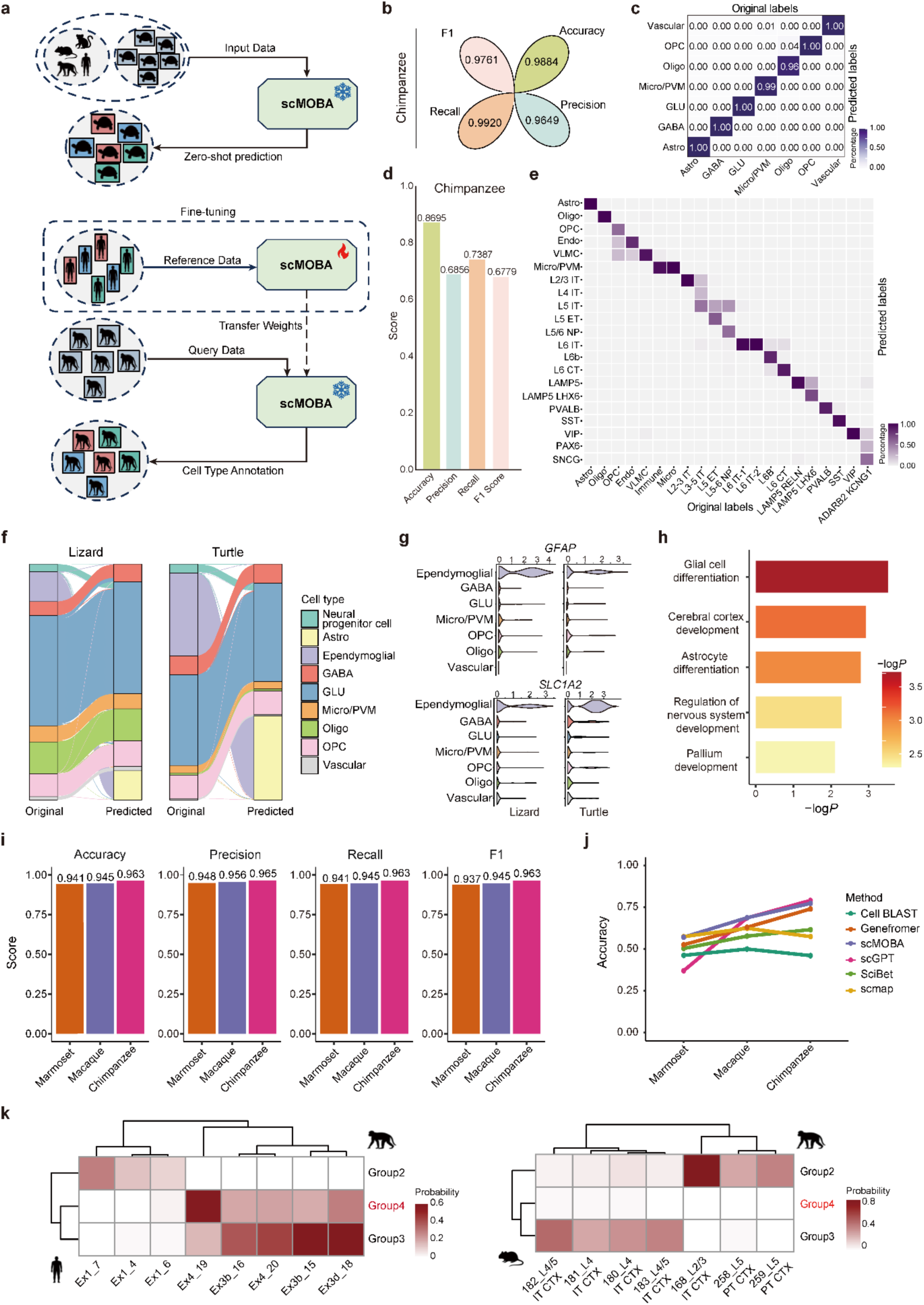
The scMOBA enables accurate cross-species cell type classification and annotation. **a**, Diagram for mapping query species data onto the referenced species data using zero-shot or fine-tuned scMOBA. **b**, Cell class classification performance on Chimpanzee snRNA-seq dataset using scMOBA zero-shot mode. **c**, Confusion matrix showing the consistency of cell class annotation generated by scMOBA and cell labels provided in the original study for chimpanzee. **d**, Bar plot showing the cell subclass classification performance on Chimpanzee snRNA-seq datasets using scMOBA zero-shot mode. **e**, Confusion matrix showing the consistency of cell subclass annotation generated by scMOBA and cell labels provided in the original study for chimpanzee. **f**, Sankey plots showing the consistency of predicted cell types by scMOBA and cell labels provided in the original study for the unseen target species lizard and turtle. **g**, Violin plots showing the expression profiles of the astrocyte marker gene *GFAP* and *SLC1A2*. **h**, Functional enrichment of top ranked genes predicted by scMOBA for the ependymoglial cells. **i**, Bar plot showing cell type classification performance of scMOBA for chimpanzee, macaque and marmoset. **j**, Comparison of cell type prediction accuracy of scMOBA and other existing methods for three non-human primate species. **k**, Heatmaps showing the prediction probabilities of glutamatergic neuron types for humans and mice using the macaque snRNA-seq dataset from the primary visual cortex (V1) as the reference.

To assess the generalization of scMOBA in evolutionarily distant species, we next applied the model to two single-cell transcriptome datasets from the brain of two reptilian species lizard and turtle^49^. The results showed that the predicted cell classes aligned well with the original annotations in both species (Fig. 4f and Extended Data Fig. 4e). The reliability of prediction was shown by the high expression of known marker gene *GAD1* in GABAergic neurons and the enrichment of GABAergic functions by top ranked genes (Extended Data Fig. 4f, g). It was noteworthy that ependymoglial cells were predicted as astrocytes, possibly because of the high similarity in gene expression profiles of these two cell types. Further examination revealed that ependymoglial cells highly expressed astrocyte marker genes including *GFAP* and *SLC1A2* (Fig. 4g). Functional enrichment showed that highly expressed genes in this cell type were involved in astrocyte differentiation and glial cell differentiation pathways (Fig. 4h), supporting its similarity with astrocytes.

We next utilized the fine-tuning mode with one species as the reference and the other as the query to further benchmark the cross-species transfer annotation ability of scMOBA (Fig. 4a). With the human dlPFC snRNA-seq dataset as the reference data, we fine-tuned scMOBA to annotate chimpanzee, macaque and marmoset cells, respectively. Consistently high (all above 0.93) performance on various metrics (accuracy, precision, recall and F1) were achieved for cell subclass prediction in all the three primate species (Fig. 4i). Despite that scMOBA showed slightly decreased performance on predicting more refined subtypes with the increasing evolutionary distance from humans, the benchmarked foundation models showed steeper performance drops than that of scMOBA (Fig. 4j).

In addition to predicting evolutionarily conserved cell types, we also studied the ability of scMOBA in identifying species-specific cell types. Specifically, we retrieved a previously published dataset comprising snRNA-seq data from cortices of humans, macaques and mice^6,50,51^. We employed a reference mapping approach to evaluate the ability to transfer cell types annotated on macaques to humans or mice (see Methods). The results showed that scMOBA accurately predicted the existence of a group of L4 glutamatergic neurons in humans but not in mice of both prefrontal cortex and V1 regions (Fig. 4k and Extended Data Fig. 4h), perfectly matching the primate-specific cell types reported in the original study^6^. These results indicated that scMOBA enables accurate transfer of cell type annotations across species and prediction of species-specific cell types.

### Accurate prediction of cellular aging with scMOBA

Given the diversity of brain corpus data used in the pretraining stage, we wondered whether scMOBA has learned the dynamic molecular changes upon complex biological processes. It is known that brain aging is complex and associated with increased risk of neurodegenerative diseases^52^. Single-cell studies have revealed that brain aging is characterized by cell-type-specific molecular shifts like pro-inflammatory glial activation and selective neuronal vulnerability^53–57^. Interestingly, we found that pre-trained cell embeddings of the scMOBA showed distinct clusters corresponding to aged individuals or young adults (Extended Data Fig. 5a), indicating the potential of scMOBA embeddings for aging clock prediction.

Therefore, we next asked scMOBA to predict the ages of cells based on the feature profiles of multi-omics data (Fig. 5a). To validate scMOBA’s capacity on biological age prediction, we retrieved a unseen snRNA-seq data from the aged human prefrontal cortex^58^, which included 7 major cell types spanning the entire life span. We fine-tuned the model with 90% of the aged human dataset to predict the ages of cells in the remaining dataset. The results showed that scMOBA predicted ages strongly correlated with the original ones, with over 99% Pearson correlation coefficients for all cell types, including astrocytes, endothelial cells, excitatory neurons, inhibitory neurons, microglia, oligodendrocytes and oligodendrocyte progenitor cells (Fig. 5b). In addition, the predicted age groups were highly consistent (over 95%) with the actual ones (Fig. 5c), and the accuracy for the predicted and actual age values also reached to over 80% for all the seven cell types (Fig. 5d). Functional analysis of top ranked genes by attention in each cell type showed consistently significant enrichment in well-known aging associated pathways including cytokine signaling in immune system, interferon signaling and pathways of neurodegeneration (Fig. 5e and Extended Data Fig. 5b-g). For examples, among the top weighted genes in endothelial cells, *HSP90AA1* was reported to promote cellular senescence and age-related tissue fibrosis by activating inflammatory and anti-proliferative signaling pathways like TGF-β and p21^59^. Another example was *ITGB1*, which is critically involved in aging by mediating cell-to-extracellular matrix adhesion and signaling^60^. Furthermore, the osteopontin encoding gene *SPP1* was ranked among the top weights in microglia (Extended Data Fig. 5d), the former was reported to contribute to neuroinflammation and neurodegeneration during aging^61–63^.

**Fig. 5.**
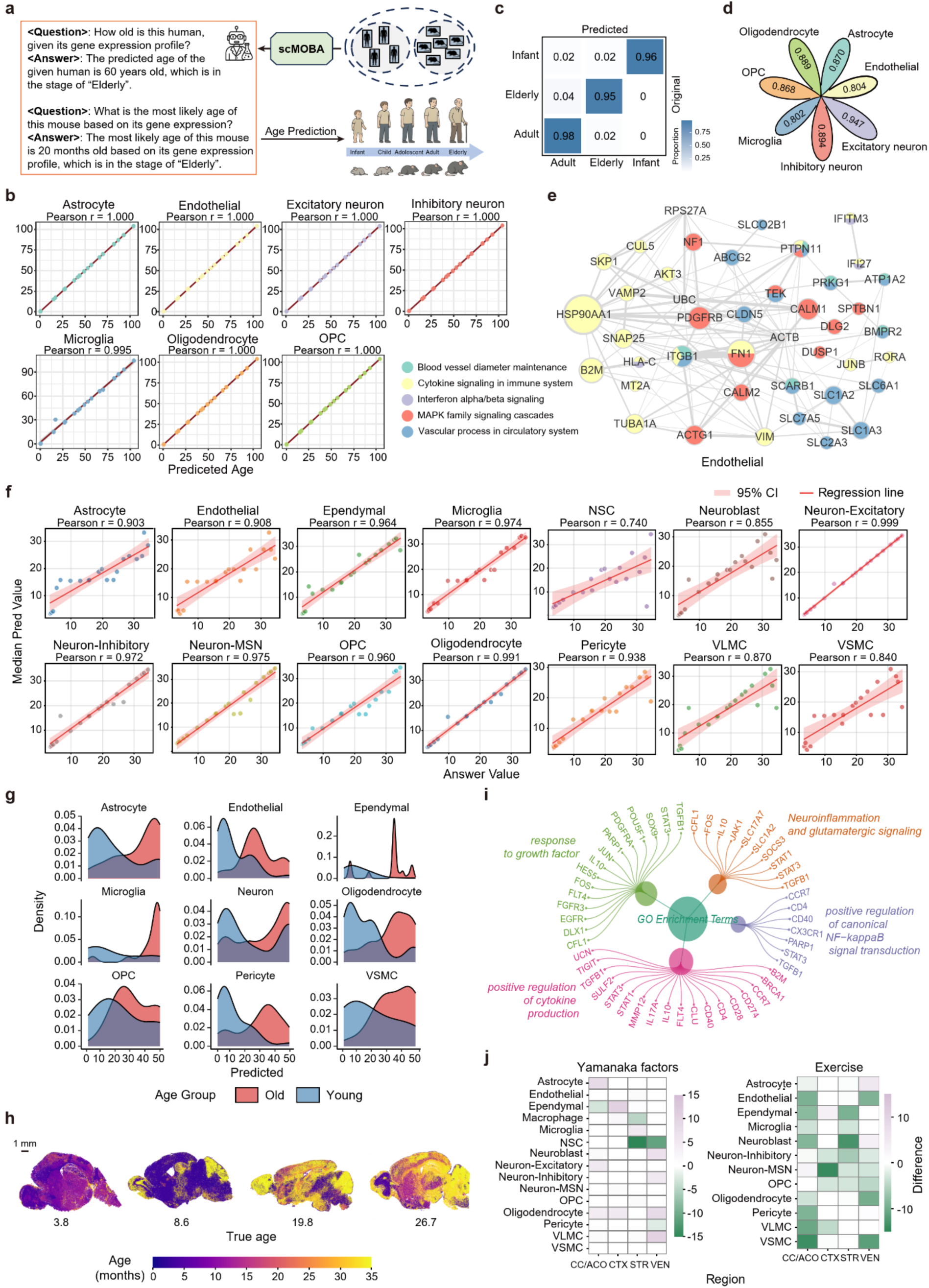
Predicting cell ages with scMOBA. **a**, Diagram showing the procedure of aging prediction by scMOBA. **b**, Dot plot showing the correlation between predicted ages by scMOBA and actual ages in the human PFC dataset^58^. **c**, Confusion matrix showing the overlap of predicted age groups by scMOBA and actual ages. **d**, Age prediction performance of scMOBA. **e**, Network plot showing the top ranked genes for predicting ages and their enriched pathways in the endothelial cells. Node colors represent associated pathways of the indicated gene. **f**, Correlation between predicted ages by scMOBA and actual ages in the mouse spatial transcriptome dataset^64^. **g**, Density plot showing the predicted age groups of scMOBA in the independent whole mouse brain spatial transcriptome dataset^64^. **h**, Spatial visualization of predicted ages in four whole mouse brain spatial transcriptomic sections with different ages. **i**, Go terms of top ranked genes for predicting ages across all cell types. **j**, Heatmaps showing the differences of scMOBA predicted ages after treatment of Yamanaka factors and exercise.

To evaluate the generality of scMOBA for aging clock prediction, we retrieved two other independent spatial transcriptome from previously published studies^61,64^. In the mouse coronal MERFISH dataset, scMOBA predicted ages showed an average of 0.92 correlation coefficient with the actual ages of mice across 14 cell types, with the highest correlation coefficient of 0.999 in excitatory neurons (Fig. 5f). In the independent sagittal MERFISH dataset comprising brain sections from different young and old mice, scMOBA also robustly distinguished old from young mice in all identified cell types (Fig. 5g). Interestingly, we mapped the predicted ages on the spatial transcriptome maps and found that the cerebellum tended to be the aging-sensitive region, since cerebella cells showed higher predicted ages than actual ones in younger ages (Fig. 5h). Similarly, functional enrichment analysis of top attention genes revealed pathways previously reported to be associated with aging, such as chronic neuroinflammation and response to growth factors (Fig. 5i), indicating the reliability of scMOBA embeddings in constructing the aging clock.

In addition, we applied scMOBA to assess the impact of two rejuvenating interventions, Yamanaka factors and Exercise on gene expression profiles of different cell types using the spatial transcriptome dataset generated from multiple regions of the aged mouse brain^64^. The scMOBA predicted that the transcriptomes of NSCs could be rejuvenated by intervention of Yamanaka factors (Fig. 5j), consistent with the result in the original study and previous report showing the restoration of neural progenitors by partial reprogramming^65^. Many cell types including endothelial, neuroblast, VLMC and VSMC in multiple brain regions were predicted to be rejuvenated by scMOBA (Fig. 5k), consistent with previous studies that exercise have strong rejuvenation effect on the brain^66,67^. Together, these results support that scMOBA embeddings have acquired meaningful representations of cell states including aging, which could be generalized to aging clock construction and prediction for independent datasets.

## Discussion

In this study, we introduce scMOBA, the first conversational agent designed to bridge the semantic gap between complex multi-omics data of the brain and human natural language. By integrating a gene encoder and a large language model through a cross-attention projector, scMOBA formulates a unified feature-question-answering framework for both unsupervised (masked gene reconstruction) and various supervised (metadata-based Q&A) tasks. This carefully designed structure alongside pretraining on the large-scale cross-species and multi-omics brain corpus data facilitated scMOBA to capture the biological processes and cellular heterogeneity underlying the complexity of brain. As a result, the model learns a consistent and information-rich objective across pre-training and downstream applications, enabling remarkable zero-shot and few-shot generalization for fine-grained cell type annotation, cross-species comparison, aging clock and disease status prediction. Furthermore, scMOBA is adaptable to customized fine-tuning strategies, enabling a range of downstream tasks such as multi-omics data integration, cell trajectory analysis, and gene regulatory network inference.

A key innovation of scMOBA lies in its pre-training strategies, which introduces unified pretraining objective and specially-designed multimodal framework that fundamentally addresses two critical limitations in the current landscape of single-cell foundation models. First, it moves beyond the paradigm of task-specific fine-tuning required by existing self-supervised models like Geneformer and scGPT. While these models excel in a “pretraining-fine-tuning” manner, their application is gated by computational expertise. Our model’s unique capability to seamlessly integrate both supervised and self-supervised pre-training paradigms allows it to learn powerful, generalizable representations that can be directly informed by biological context. This mitigates the reliance on extensive fine-tuning for every new downstream task, thereby lowering the barrier for non-computational neuroscientists.

Second, our model transcends the narrow focus of existing approaches. While recent efforts have begun to leverage LLMs for specific tasks like cell type annotation, they are often constrained by their reliance on textual knowledge alone^31,68,69^, leading to concerns about hallucination and an inability to capture the dynamic, context-dependent nature of biological systems. LangCell^70^ and CellWhisper^71^ attempted to connect transcriptomes with text for interactive exploration of single-cell RNA-sequencing data by contrastive learning, which underlie the chat-based transcriptome analysis. Nevertheless, our multimodal architecture, inspired by VLMs, is not a single-task tool. By aligning multi-omics feature abundance profiles with natural language, we create a flexible framework capable of a broad range of inference tasks. Moreover, the learned generalizable representations and their compatibility with customized fine-tuning strategies ensures that the model is not limited to cell type annotation but can be readily personalized and extended to complex downstream analyses such as multi-omics integration, temporal analysis, and gene regulatory network inference. Therefore, our model serves not merely as a predictor but as a versatile biological expert system, greatly bridging the gap between powerful foundational AI and practical, accessible scientific discovery.

Another merit of scMOBA is its flexible and efficient training scheme. The two-stage pre-training process - beginning with cross-modal alignment via a trainable projector, followed by joint optimization with LoRA-based LLM fine-tuning - ensures that the model retains the representational power of both the gene encoder and the LLM. This hierarchical training strategy allows scMOBA to reason over complex multi-omics contexts without catastrophic forgetting or overfitting. The novel pre-training strategies of scMOBA increase its ability to scale across diverse biological dimensions through a unified feature-text representation, enabling pre-training on a wide range of biological data, including various omics types, species, and biological states. By leveraging this scalable architecture, scMOBA can seamlessly integrate and generalize across a variety of single-cell datasets and biological contexts, improving its adaptability and robustness in tackling novel research datasets and tasks.

The model’s ability to accurately predict cell types across independent snRNA-seq, snATAC-seq, and spatial transcriptomic datasets - spanning humans, non-human primates, and even evolutionarily distant reptilian species - underscores its robust cross-species and cross-modality capabilities. Notably, scMOBA outperforms existing benchmarks in both fine-grained cell type annotation and integration of heterogenous data modalities, achieving superior batch removal and biological conservation scores. Furthermore, scMOBA enables the identification of evolutionarily conserved and species-specific cell types, as evidenced by its accurate transfer of annotations across primates and the detection of a primate-specific L4 glutamatergic neuron subclass. This suggests that the embeddings learned by scMOBA capture fundamental biological signals while effectively mitigating technical variations.

Looking forward, the conversational interface of scMOBA opens new avenues for interactive and hypothesis-driven single-cell analysis. Researchers can query the model in natural language to perform tasks ranging from cell type annotation and spatial mapping to regulatory network inference. As the scale and diversity of single-cell datasets continue to grow, models like scMOBA will play an increasingly critical role in unifying multi-omics data beyond transcriptomics and epigenomics (e.g., proteomics and metabolomics) into a biologically meaningful and accessible framework. Additionally, the structure of scMOBA was also extensible to other tissues or the whole organisms for integration and exploration.

In conclusion, scMOBA represents a pivotal step toward a more integrated, interpretable, and scalable paradigm for constructing “digital brain”. By harmonizing feature representations of multi-omics data with natural language reasoning, it not only facilitates accurate and efficient biological discovery but also lays the foundation for future AI-driven systems capable of contextualizing cellular function across development, evolution, aging, and disease.

## Acknowledgements

We thank for the support of Shanghai Artificial Intelligence Laboratory. This work was conducted during the internships of Ran Wei, Jianle Sun, Peng Zheng, Chaoqi Liang, and Fanyi Meng at the Shanghai Artificial Intelligence Laboratory. The project was supported by National Science and Technology Innovation 2030 Major Program (STI2030-2021ZD0200100), National Key Research and Development Program of China (No. 2024YFC3408000), National Natural Science Foundation of China (NSFC) Outstanding Youth Foundation (No. T2422026), National Natural Science Foundation of China (No. U23A6010) and Shanghai Science and Technology Development Funds (23QA1410400).

## Author contributions

Methodology, R.W., Z.Z., P.Y. and Y-D.S.; Investigation, R.W., Z.Z., J.S. and Y-K.S.; Formal Analysis, R.W., Z.Z., J.S., Y-K.S., J.M. P.Z., C.L. and F.M.; Writing – Original Draft, Y-D.S.; Writing – Review & Editing, R.W., Z.Z., J.S., Y-K.S., J.M., P.Z., C.L., F.M., P.Y. and W.O.; Conceptualization and supervision, L.B., P.Y. and Y-D.S..

## Methods

### Datasets and processing

#### Datasets

##### snRNA-seq datasets and preprocessing

The snRNA-seq datasets were compiled from multiple public resources, including the CELLxGENE Portal, the Gene Expression Omnibus (GEO) repository, the Human Cell Atlas (HCA), the Brain Cell Atlas (BCA), the Allen Brain Cell Atlas, and published studies (Supplementary Table 1). Collectively, these datasets encompass 2.69 million brain cells across four species: mouse (Mus musculus), marmoset (Callithrix jacchus), crab-eating macaque (Macaca fascicularis), and human (Homo sapiens).

We downloaded the processed data, manually converted all datasets into the h5ad format to ensure compatibility with our Python-based analysis pipeline and further transformed the raw count matrices into normalized expression values. We used ‘sc.pp.normalize_total’ and ‘sc.pp.log1p’ function from scanpy with parameters set to target_sum= 10000, ultimately generating a matrix with normalized expression values.

#### snATAC-seq datasets and preprocessing

##### Alzheimer’s disease multi-omics snATAC-seq

The snATAC-seq data were obtained from the GSE174367 dataset^74^ accessible from the NCBI Gene Expression Omnibus (GEO) database. The dataset profiles chromatin accessibility in 130K nuclei from postmortem human brain tissues of individuals with late-stage Alzheimer’s disease and healthy controls, enabling the identification of cell-type-specific regulatory alterations in disease pathogenesis. We downloaded the processed data, which included the filtered peak-barcode matrix and cell metadata.

##### Human brain scATAC-seq atlas for neurodegenerative diseases

We retrieved scATAC-seq data from the epigenomic atlas GSE147672 (NCBI GEO)^75^, which characterized 42K cells from six distinct brain regions of 39 cognitively normal individuals. This comprehensive atlas maps 359,022 cell-type-specific regulatory elements across 24 brain cell clusters and provides a healthy reference framework for identifying genetic risk loci for Alzheimer’s and Parkinson’s diseases through its integrated multi-omics approach.

##### Mouse primary motor cortex

The snATAC-seq data from the mouse primary motor cortex (MOp) cell atlas was accessed from the BRAIN Initiative Cell Census Network (BICCN)^46^. This dataset provides a high-resolution epigenomic profile of the region crucial for motor control. We utilized the processed snATAC-seq data, which includes the cell-by-bin matrix and cell-type annotations. For the purpose of our analysis, we focused on the snATAC-seq modality to investigate the cis-regulatory landscape. The final dataset contained 71K nuclei. During data processing, we followed the original BICCN processing pipeline and retained the consensus set of regulatory elements for analysis.

We used the ‘GeneActivity’ function from Signac with parameters set to extend.upstream = 2000, extend.downstream = 200, and max.width = 500000, ultimately generating a matrix with genes as rows and cells as columns. This matrix represents the gene activity scores.

##### Spatial transcriptome and preprocessing

The spatial transcriptomics data for human were primarily sourced from previous studies listed in Supplementary Table 1. Mouse data were mainly obtained from the Allen Brain Atlas, while data for marmoset and macaque were predominantly derived from the Non-Human Primate Brain Atlas established by the Center for Excellence in Brain Science and Intelligence Technology, Chinese Academy of Sciences. We downloaded the processed spatial transcriptomics datasets and applied the same preprocessing pipeline used for the snRNA-seq data.

##### Alignment of cell metadata across modalities and species

Due to scMOBA’s novel input design, we curated five key metadata fields in the .obs attribute: ‘Class’, ‘Subclass’, ‘Tissue’, ‘Developmental stage’, and ‘Disease’. Among these, ‘Class’ and ‘Subclass’ denote the broad and fine-grained cell types, respectively. ‘Tissue’ indicates the brain region from which the cell was sampled. ‘Developmental stage’ reflects the donor’s age or developmental stage, and ‘Disease’ specifies whether the donor was affected by a disease and, if so, the specific diagnosis.

##### Consensus cell types

We standardized the nomenclature of cell types in the ‘Class’ and ‘Subclass’ annotations. Across all four species, we harmonized the ‘Class’-level cell types into the following categories: GLU (glutamatergic neurons), GABA (GABAergic neurons), Astro (astrocytes), Oligo (oligodendrocytes), OPC (oligodendrocyte precursor cells), Micro/PVM (microglia and perivascular macrophages), Ependymal, Vascular, Choroid Plexus, Fibroblast, and Bergmann glia. However, due to the inherent challenges in aligning ‘Subclass’-level cell types across species, we performed subclass harmonization only within each individual species. A detailed list of subclass annotations is provided in the Supplementary Table 2.

##### Developmental stage

For the gestational stage, we adopted the convention of “GW + number” for human samples and “E + number” for mouse, marmoset, and macaque samples, consistent with established usage in the field. For post-gestational stages, we used “number-week-old stage” or “number-year-old stage” for human samples and “P + number” for mouse, marmoset, and macaque samples.

##### Brain regions

We retained the original brain region annotations of the samples and allowed the LLM component of our model to automatically process this information.

##### Disease status

We applied the same preprocessing pipeline as used for ‘Brain regions’ but specifically unified all healthy samples under the label ‘Normal’.

### Pre-processing for the scMOBA

We assembled a large-scale cross-species multi-omics pretraining corpus, comprising 38.3 million human cells (including 18.8 million snRNA-seq, 1.5 million snATAC-seq and 18 million spatial transcriptome measurements), 30.2 million mouse cells (including 10.9 million snRNA-seq, 2.3 million snATAC-seq, and 17 million spatial transcriptome measurements), 48.2 million macaque cells (including 4.6 million snRNA-seq, 1.6 million snATAC-seq, and 42 million spatial transcriptome measurements), and 12.9 million marmoset cells (including 0.5 million snRNA-seq, 0.4 million snATAC-seq, and 12 million spatial transcriptome measurements).

Different data sources, omics layers and labels are harmonized (see Supplementary Materials). We established the correspondence between gene names and Ensemble IDs within each dataset, and converted orthologous genes from Ensemble across different species to a unified ID using a homology table (Supplementary Table 3). For other non-RNA omics data, feature quantification was uniformly mapped to the gene scale, such as converting snATAC-seq peak accessibility signals into gene activity scores.

We adopted the rank-value tokenization method from Geneformer, which ranks genes based on their expression within the cell and normalizes these values against their expression across the entire pre-training corpus for each specie and omics. This approach is designed to highlight marker genes and highly variable genes (HVGs) that are actively expressed and reflective of cellular differences. Specifically, we calculated the non-zero median expression for each detected gene across the corpus. Genes were then normalized using this non-zero median, and ranked based on their normalized expression level within that specific cell. Finally, the rank-value encoding for each single-cell transcriptome was tokenized using a master vocabulary constructed across all datasets within each omics and each specie. This vocabulary also includes two special tokens used for padding and masking.

It is noteworthy that we employed a uniform tokenization strategy to construct distinct vocabularies for datasets derived from different species and omics platforms. This consistent methodology was crucial, as it permitted the use of a unified encoder for model training, thereby allowing the model to simultaneously learn specific features from each species and omics layer.

### scMOBA Architecture and Pretraining Framework

#### scMOBA architecture

scMOBA is a multi-modal architecture designed to achieve deep integration between feature abundance data and textual information. The motivation for adopting a multi-modal framework lies in the complementary nature of biological and linguistic modalities: while numerical feature abundance profiles capture quantitative cellular states, textual data encode rich biological knowledge and interpretability. By combining these two sources of information, scMOBA enables comprehensive understanding and reasoning across both molecular and semantic spaces, thereby enhancing interpretability and generalization in single-cell biology.

The architecture consists of three major components: a gene encoder, a projector, and a large language model. The gene encoder comprises six Transformer encoder units, each containing a self-attention layer and a feed-forward neural network layer. It takes an input size of 2,048 with an embedding dimension of 256 and four attention heads per layer. Initialized with pretrained weights from Geneformer, the encoder inherits biologically meaningful representations learned from large-scale single-cell corpora. Owing to the flexibility of the Transformer structure, this encoder can be readily extended to handle multi-omics modalities (e.g., transcriptomic, epigenomic, and spatial data) and cross-species datasets, facilitating transfer learning across diverse biological systems.

The projector module includes two graph convolutional network (GCN) layers, which further model gene–gene interactions based on the encoded feature representations. To effectively summarize informative biological signals, 256 learnable queries are introduced to perform cross-attention with the GCN outputs, extracting the most relevant features for downstream textual reasoning while reducing the input length to the language model. Finally, the language model component—built upon a 2-billion-parameter LLaMA architecture—receives the concatenated gene-derived features and text-formatted queries as input. It jointly processes these signals to generate natural language outputs, enabling end-to-end reasoning across molecular and linguistic modalities. Collectively, this design allows scMOBA to perform biologically grounded interpretation and knowledge synthesis in a unified multimodal framework.

#### Construction of question–answer pairs

To align the pretraining process with diverse downstream tasks, scMOBA adopts a question–answer –based pretraining strategy. This design allows the model to directly learn in a task format consistent with its eventual application, such as biological reasoning, annotation, or interpretation in natural language. By framing biological knowledge in a QA form, the model effectively integrates supervised signals—derived from curated metadata and cell annotations—with unsupervised information learned from large-scale gene expression patterns. Moreover, this format facilitates generalization across multiple species and multi-omics datasets, since biological relationships and descriptive attributes can be expressed in a unified textual representation, regardless of data modality or organismal origin.

For the construction of question–answer pairs, each cell in the dataset is accompanied by metadata providing supervisory information such as cell type, subclass, age or developmental stage, and disease state. These metadata attributes are used to formulate text-based questions paired with corresponding answers, enabling the model to learn biologically grounded associations between gene expression and descriptive annotations.

To enhance the model’s understanding of gene–gene relationships, a subset of genes within each cell is randomly masked, and the model is trained to predict their expression values. In addition, to capture spatial dependencies, we design a spatial prediction task in which the expression profiles of neighboring spots are provided as context, and the model predicts highly expressed genes in the central cell.

To further enhance linguistic diversity, multiple templates are designed for each type of question–answer pair. During data construction, one template is randomly selected for each instance to ensure variation in both the textual input and output of the language model. Through this integrated pretraining framework, both supervised biological annotations and unsupervised gene expression patterns are jointly leveraged, allowing scMOBA to unify multimodal biological knowledge within a single learning paradigm.

#### Learning objective for pretraining

The objective of scMOBA pretraining is to generate accurate textual responses conditioned on both the gene expression input and the text-formatted question.

During pretraining, the model is trained in an autoregressive manner, where the large language model predicts the next token *t_i_* given the preceding tokens and multimodal context. The overall loss function is defined as the standard language modeling loss over all tokens in the answer sequence:

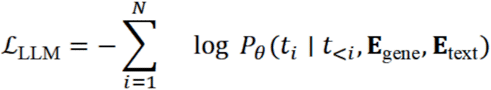

where **E**_gene_ and **E**_text_ represent the embeddings of gene expression and textual input, respectively, and *P_θ_* denotes the model’s predicted probability distribution parameterized by *θ*.

This objective encourages the model to learn joint representations across the gene and text modalities while maintaining the generative capabilities of the language model. By optimizing the autoregressive likelihood, scMOBA effectively aligns biological signals with semantic information, enabling coherent biological question answering during inference.

#### Two-stage pretraining strategy

scMOBA adopts a two-stage pretraining strategy to progressively align gene expression representations with the language model’s semantic space while preserving the pretrained capabilities of each modality.

In the first stage, the parameters of both the gene encoder and the language model are frozen. The gene encoder is initialized from Geneformer, which already possesses a strong ability to capture gene expression representations, while the language model retains its intrinsic linguistic competence. During this stage, only the projector parameters are optimized. The projector learns to map the latent representations from the gene encoder to the semantic embedding space of the language model, achieving modality alignment through cross-attention and contrastive supervision. This stage focuses on bridging the representational gap between the gene and text modalities without disturbing their pretrained knowledge.

In the second stage, all components are unfrozen for joint training. The large language model is fine-tuned with LoRA (Low-Rank Adaptation) to preserve its general language ability while adapting to biological reasoning tasks. The full model—including the gene encoder, projector, and LoRA adapters in the LLM—is optimized end-to-end using the autoregressive language modeling objective. This stage allows deeper multimodal fusion, enabling the model to generate biologically consistent and context-aware textual responses from gene expression inputs.

#### Downstream tasks

Downstream analyses were formulated in the unified gene–question–answer format, allowing the pretrained scMOBA model to perform diverse biological tasks without task-specific retraining. Below we describe the major categories of downstream evaluations.

#### Cell type classification

scMOBA was evaluated on multiple single-cell transcriptomic datasets for fine-grained cell-type classification across species and modalities. For each cell, the model received its gene-expression (or ATAC-derived gene activity) vector and a textual question such as *“What is the cell type of this cell?”*. In the zero-shot setting, the model directly generated the predicted cell type from the pretrained parameters. In the fine-tuned setting, scMOBA was evaluated under both a small dataset and supertype classification schemes. In the small dataset mode, the model was trained on a small subset of annotated examples to refine the projector and LLM modules. Subclass-level classification was performed on human, mouse, and marmoset datasets, using 20% of the data for training and 80% for evaluation. Model performance was assessed using accuracy, precision, recall, and F1 score. For supertype classification, 80% of the data were used for training and 20% for testing.

#### Integration task

The integration task aimed to align data from different omics, species, and experimental batches into a unified embedding space. Because integration depends on dataset-specific characteristics, fine-tuning was required for each configuration of species, modality, and batch. During fine-tuning, scMOBA was optimized with two complementary loss components. The main loss (L_main) was the language modeling loss, implemented as a next-token prediction objective that trained the model to generate accurate textual answers (e.g., predicted cell types) from multi-omics inputs. In parallel, a domain loss (L_domain) was introduced as a domain classification objective, encouraging the model to identify the origin of each sample (e.g., species, modality, or batch). Gradients from the domain classifier were reversed during backpropagation to promote domain-invariant representations and suppress dataset-specific signals. The total training objective was defined as:

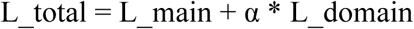

where α = 0.5 by default but could be adjusted according to dataset complexity. This joint objective enables the model to maintain strong language-driven prediction ability while progressively minimizing domain-specific biases, resulting in effective and biologically meaningful multi-omics integration.

Integration performance was evaluated using metrics from the scIB benchmark, including KMeans-NMI, KMeans-ARI, Silhouette-label, Graph-connectivity, and Silhouette-batch scores. The first three metrics measure the preservation of biological structure and cell-type separability, whereas the latter two assess the degree of domain removal and batch correction quality.

#### Cross-species transfer analysis

For cross-species integration, the experimental setup followed the same configuration as the multi-omics integration task. In the cross-species comparison experiments, the pretrained scMOBA model was directly applied to datasets from chimpanzee, rhesus macaque, lizard, and turtle without additional fine-tuning. For the comparisons among human, macaque, chimpanzee, and marmoset, subclass- and supertype-level classification was performed by training on human data and testing on the other three species to evaluate the consistency of cell-type representation across species. To assess species specificity, the model was trained on macaque cells and evaluated on human and mouse datasets, enabling the identification of species-specific and conserved transcriptional patterns.

#### Aging clock construction

We constructed aging clocks for both human single-cell transcriptomic data and mouse spatial MERFISH datasets to predict biological age. For the human snRNA-seq dataset, we randomly split the data into 90% for training and 10% for testing. For the mouse MERFISH data, we applied spatial smoothing to denoise local transcriptomic variations prior to age prediction, and used a 90/10 split on the coronal dataset for model training and testing. In all settings, a single unified model was trained to predict the biological age of each cell type. The model performance was assessed by computing Pearson correlation coefficients between the predicted and chronological ages in the held-out test sets. To further examine cross-region generalization, the mouse aging clock trained on the coronal MERFISH data was additionally evaluated on an independent sagittal dataset, enabling assessment of its robustness to spatial and anatomical shifts. To validate the biological relevance of the learned aging signatures, the trained aging clocks were further tested under two rejuvenation paradigms—exercise intervention and Yamanaka factor induction—to determine whether the models could capture rejuvenation or aging-reversal effects. Together, these analyses demonstrate the strong predictive power, cross-dataset generalization ability, and biological interpretability of scMOBA-based aging clocks.

#### Gene-level attention score computation

To identify genes most attended by the model, we quantified gene attention scores using a two-stage attention aggregation. For each sample, we first extracted the LLM first-layer attention between the final question token and each learnable query vector. Let

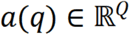

denote these query-level attention weights (averaged over heads).

Next, we computed cross-attention between each learnable query and all GCN-derived gene tokens. For query i, the attention over gene tokens is

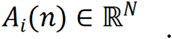

The final attention score for each gene token jjj was obtained by a weighted sum over all queries:

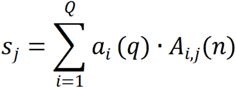

Scores were averaged across the batch dimension to obtain

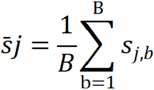

Each token position was mapped to its corresponding gene ID using the Geneformer token dictionary, and gene-level scores were aggregated across the entire test set. We further normalized scores by the number of samples and applied min–max scaling to derive final gene attention values.

#### Protein interaction analysis

For each cell type, we selected the top 300 genes ranked by gene attention weights, and then predicted the protein-protein interaction network using STRING (v.12.0) database with default parameters. GO enrichment analysis was further performed by Metascape (https://metascape.org/). Finally, subnetworks were presented with Cytoscape (v.3.10.1) tools.

### Benchmarking experiment setup

#### Cell type classification

To evaluate the cross-species cell annotation utility of foundation models, we benchmarked Geneformer and scGPT against each other. We performed fine-tuning using subsets of data from human, marmoset, and mouse datasets—specifically those unseen during pre-training—and assessed cell subclass prediction on the held-out samples. Performance was measured using accuracy, macro-precision, recall, and F1 scores. Furthermore, to characterize label efficiency (dependency on labeled data), we systematically evaluated model performance across training splits ranging from 5% to 50% with a step size of 5%.

We further assessed the model’s ability to predict more granular cell super-types using human datasets. Models were fine-tuned on an 80% data split and evaluated on the remaining 20%

#### Integration task

In this work, we performed a comprehensive benchmark of five integration methods—CCA Integration, RPCA Integration, Harmony Integration, FastMNN Integration (all implemented using Seurat v5), and scVI—for integration tasks.

Four of the five methods (CCA, RPCA, Harmony, and FastMNN) shared a consistent preprocessing pipeline where gene expression values were normalized per cell based on total counts and subsequently log-transformed. The same set of 2,000 highly variable genes (HVGs) was selected for all methods; however, for scVI, we provided the unnormalized count data of these HVGs instead of the processed values used for the other methods.

For integration analysis, all methods were configured to generate 50-dimensional embeddings. The four Seurat-based methods were performed using the IntegrateLayers function with their respective algorithms, while scVI was implemented through its standard generative modeling framework to obtain 50-dimensional latent representations. This consistent dimensionality enabled direct comparison of integration performance across the different methods.

Comparisons were also made with existing foundation models. In accordance with the official tutorials, Geneformer’s integration performance was evaluated using zero-shot cell embeddings, while scGPT was fine-tuned using batch and cell type labels. All benchmarking was performed using scaled scIB metrics.

#### Cross-species transfer analysis

We conducted a systematic benchmark of three cell type annotation methods—scmap, SciBet, and Cell BLAST—for cross-species cell type annotation tasks. Both scmap and SciBet adopted a log-transformation-based preprocessing pipeline. Specifically, scmap applies log transformation after adding a pseudo-count of 1 to the raw count data, while SciBet performs library size normalization using the CPM function from the edgeR package. Both methods uniformly selected 2000 highly variable genes (HVGs): scmap achieved this through its built-in selectFeatures function, and SciBet used the ‘SelectGene_R’ function from its package for gene selection. In contrast, Cell BLAST directly uses raw count data as input and identifies HVGs via the ‘find_variable_genes’ function with default parameters. The union of HVGs from both the reference and query datasets was used for downstream analysis.In the cell type annotation step, all three methods generated cell type predictions based on their respective algorithmic frameworks. scmap and SciBet performed annotation using their core functions—’scmapCluster’ with a threshold set to 0.1 and ‘SciBet_R’, respectively—while Cell BLAST applied batch effect correction through its ‘fit_DIRECTi’ function with lambad_reg=0.001. This standardized evaluation framework ensures direct comparability of performance across different annotation methods.

## Supplementary figures

**Extended Data Fig. 1.**
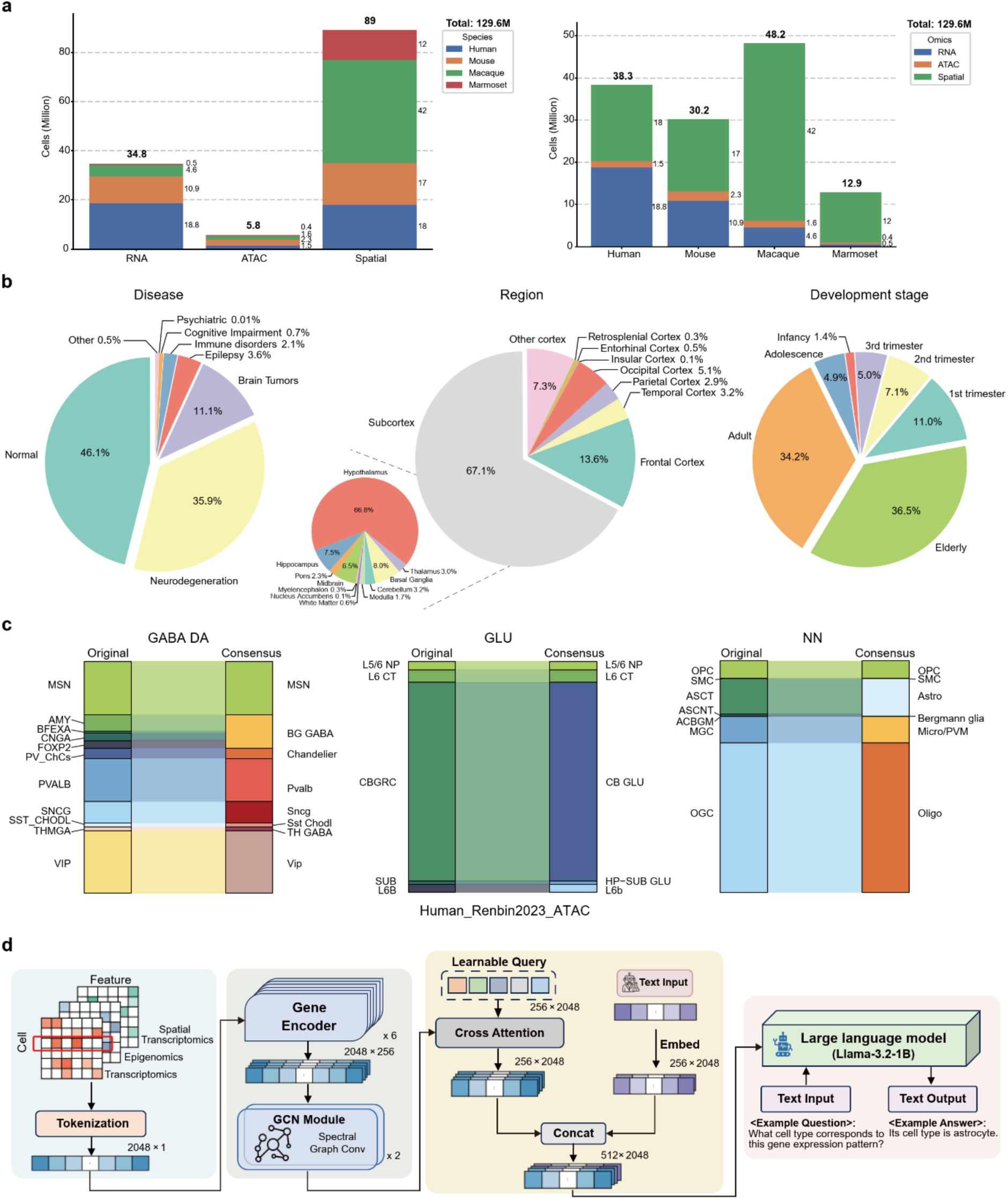
Summary of the datasets and scMOBA architecture. **a**, Numbers of cells from different modalities and species used in this study. **b**, Pie charts showing the proportions of cell origins, including diseases, brain regions and developmental stages. **c**, Sankey plot showing the consistency of consensus cell labels used in the pre-training datasets and those provided in the original studies. **d**, Detailed architecture of scMOBA.

**Extended Data Fig. 2.**
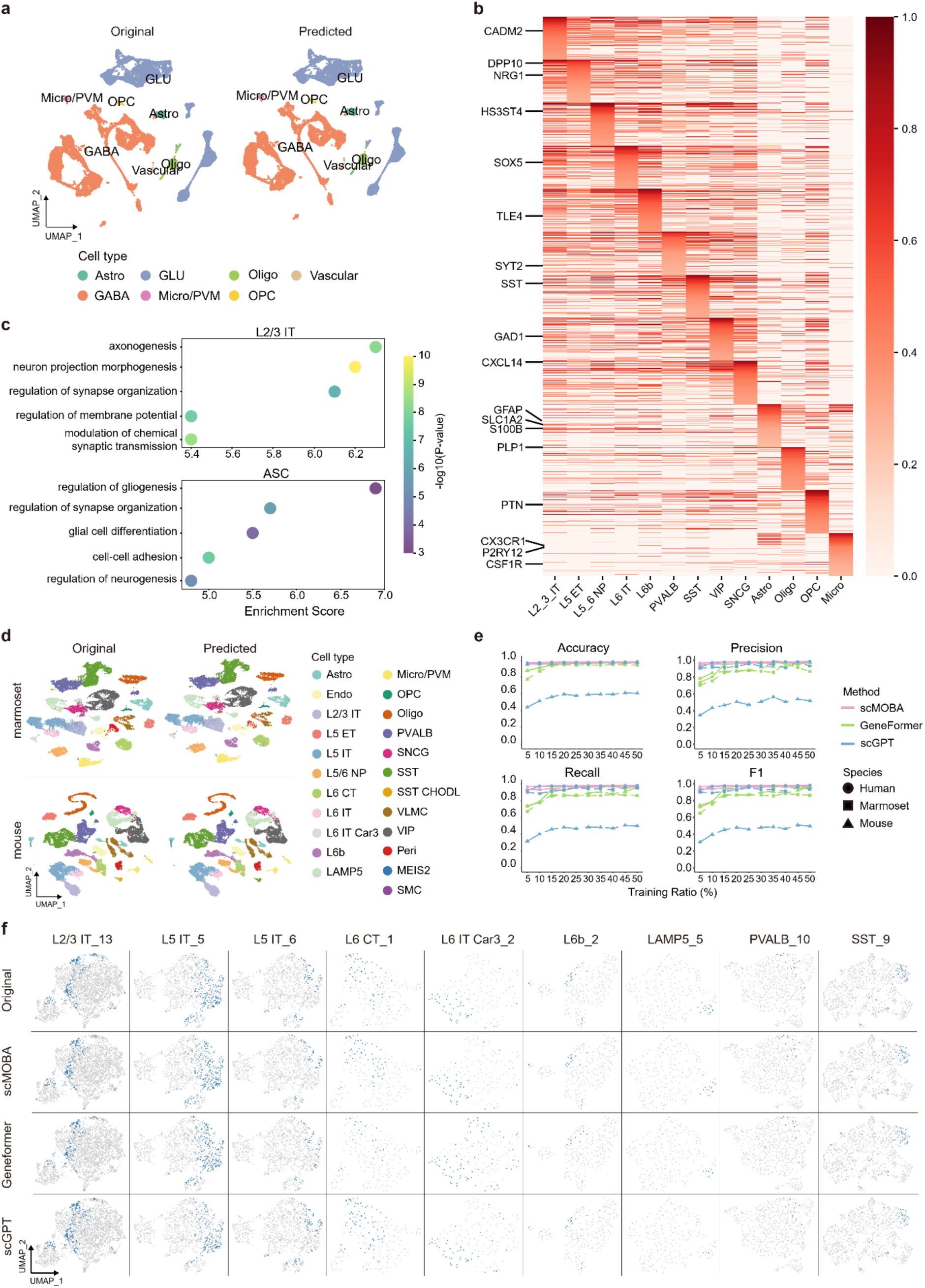
Cell type-annotation results using scMOBA. **a,** UMAP plots of gene expression profiles from the M1 snRNA-seq dataset, colored by cell classes provided in the original study (left) and those predicted by scMOBA (right). **b**, Heatmap showing the normalized weights of top 200 ranked genes for each cell subclass. **c**, Functional enrichment analysis of top 200 ranked genes for L2/3 IT and Astrocytes (ASCs). **d**, UMAP plots of gene expression profiles from the M1 snRNA-seq dataset, colored by cell subclasses predicted by Geneformer and scGPT. **e**, Prediction accuracy of scMOBA, Geneformer and scGPT in cell subclass annotation using different training-validation split cut-offs for the few-shot training strategy. **f**, UMAP visualizations of predicted cell clusters by scMOBA, scGPT and Geneformer and cell labels provided by the original study^73^.

**Extended Data Fig. 3.**
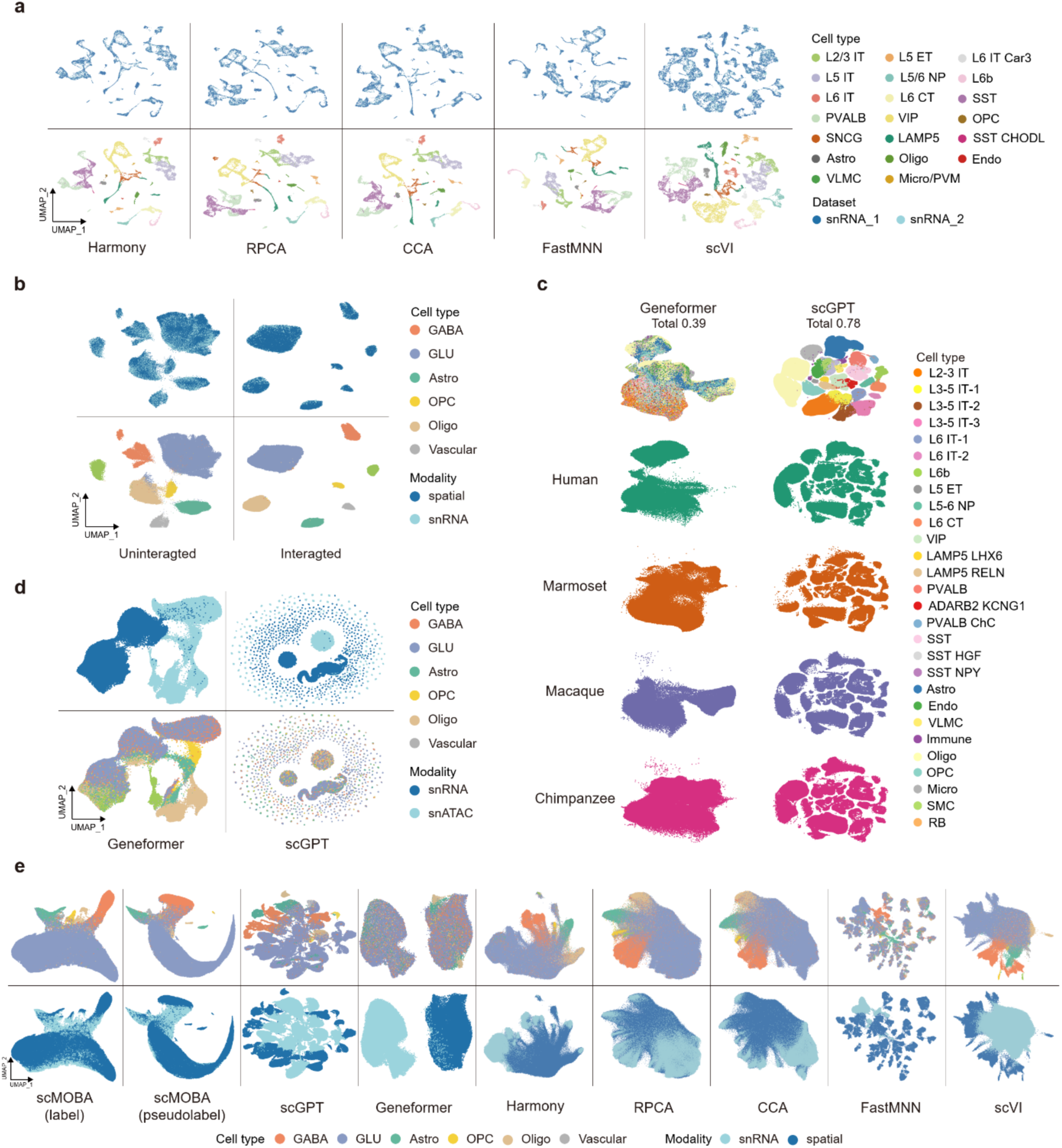
Multi-omics and cross-species integration using scMOBA. **a**, UMAP plots showing the integration results of two batched snRNA-seq datasets using cell embeddings of Harmony, Seurat v4 RPCA, Seurat V3 CCA, fastMNN, and scVI. Cells are labeled by their cell types. **b**, UMAP visualization for cell embeddings of two spatial transcriptome slides before (left) and after (right) scMOBA integration. **c**, UMAP visualization of snRNA-seq data from four primate species using cell embeddings of different integration methods. **d**, UMAP visualization of snRNA-seq and snATAC-seq data integration results using Geneformer and scGPT. **e**, UMAP visualization of cell embeddings of snRNA-seq and spatial transcriptome data of the macaque cortex. Cells are colored by cell subclasses

**Extended Data Fig. 4.**
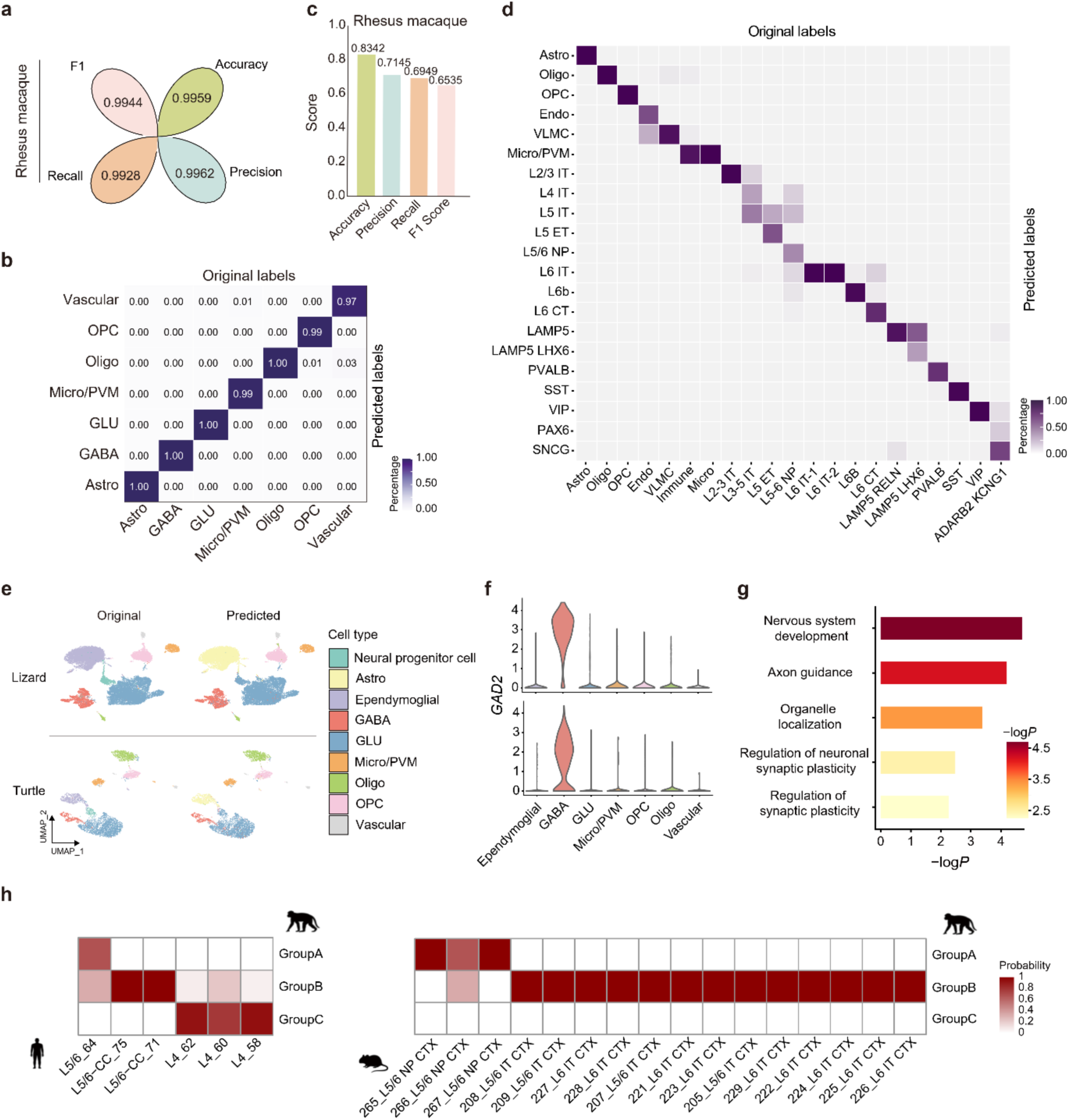
Benchmarking performance of scMOBA for cross-species cell-type annotation. **a**, Cell class classification performance on Macaque snRNA-seq datasets using the zero-shot mode of scMOBA. **b**, Confusion matrix showing the consistency of cell class annotation generated by scMOBA and cell labels provided in the original study for Rhesus macaque. **c**, Bar plot showing the cell subclass classification performance on Rhesus macaque snRNA-seq dataset using scMOBA zero-shot mode. **d**, Confusion matrix showing the consistency of cell subclass annotation generated by scMOBA and cell labels provided in the original study for Rhesus macaque. **e**, UMAP plots of cell types predicted by scMOBA and those provided in the original study for the lizard and turtle snRNA-seq datasets. **f**, Violin plots showing the expression profiles of the GABAergic marker gene *GAD2*. **g**, Functional enrichment of top ranked genes predicted by scMOBA for GABAergic cells. **h**, Heatmaps showing the prediction probabilities of glutamatergic neuron types for humans and mice using the macaque snRNA-seq dataset from the primary frontal cortex (PFC) as the reference.

**Extended Data Fig. 5.**
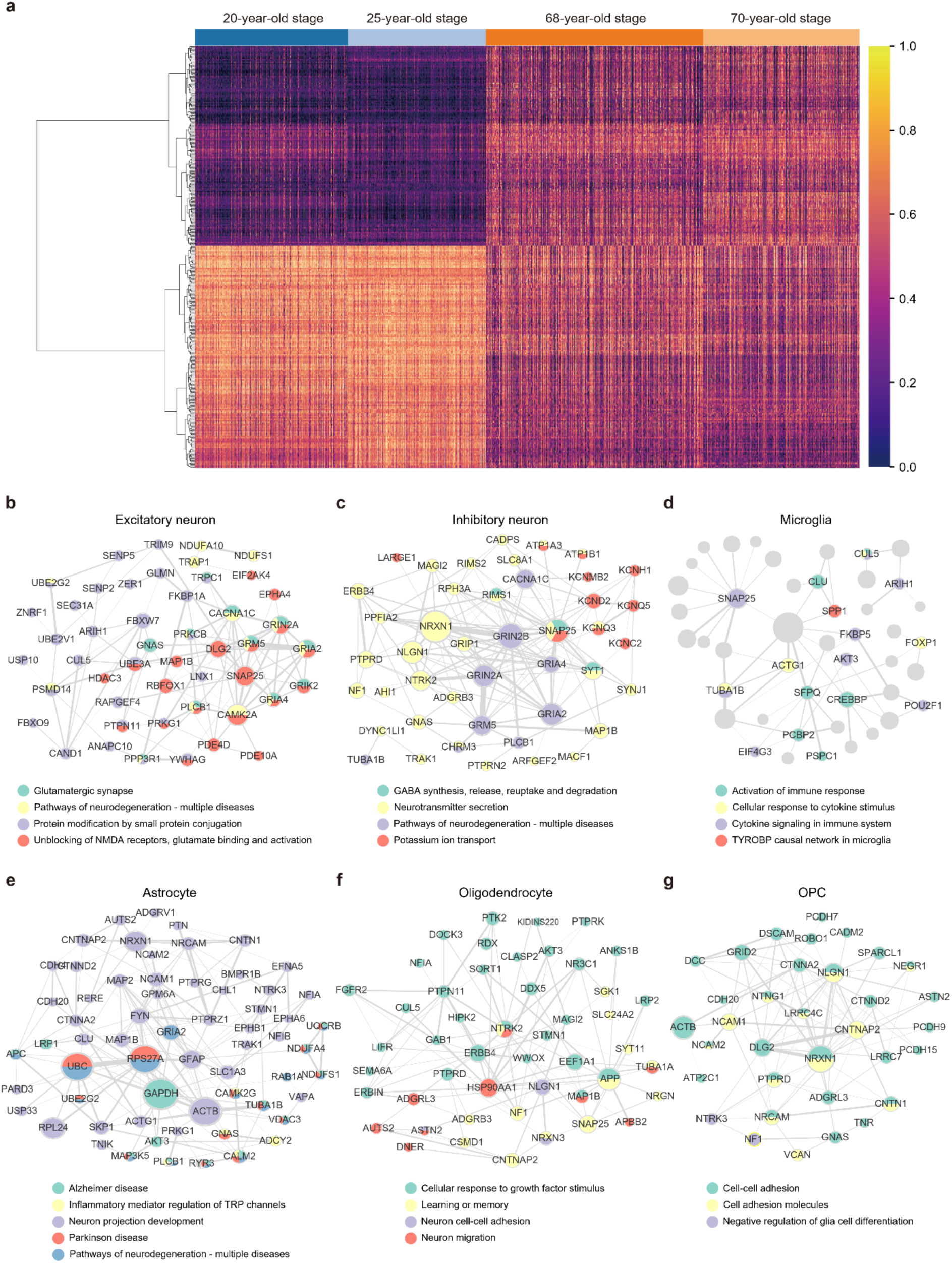
Cell embeddings of scMOBA are distinguishable for different age groups. **a**, Heatmap showing the cell embeddings of scMOBA for different age groups. **b-g,** Network plots showing the top ranked genes for predicting ages and their enriched pathways in the excitatory neurons (b), inhibitory neurons (c), microglia (d), astrocyte (e), oligodendrocyte (f) and OPC (g).

